# Linguistic structure reshapes neural representations of word meaning

**DOI:** 10.64898/2026.09.23.753793

**Authors:** Simone M. Krogh, Liina Pylkkänen

**Author notes:** **CORRESPONDING AUTHOR:** Simone M. Krogh, 10 Washington Place, New York University, New York, NY 10003, United States.

## Abstract

In language, not only do sentence meanings depend on how words are structured together, but word meanings also depend on their roles in the structure. While we have some understanding of how the brain grasps complex meanings built from simpler parts, we know far less about how linguistic structure reshapes the meanings of the words themselves. Here we used magnetoencephalography to track the neural dynamics of animacy, a fundamental semantic feature that guides conceptual organization. Regardless of how animacy was operationalized (as discrete semantic categories or more abstractly), the same pattern emerged: When word-level animacy conflicted with the structural context, it disappeared from the word’s neural representation. Our findings show how compositional processes rapidly reconfigure neural traces of word meanings, allowing phrase-level meanings to even override word-level features.

**SIGNIFICANCE STATEMENT:** When we use the word *tiger,* it evokes the nonlinguistic concept of a tiger and allows us to convey the meaning of a wild striped feline. Here, we asked whether the neural signatures of words are stable when used across different structural positions: Is *tiger* the same animal in *valley tiger* and *tiger valley*? We found that phrase-level animacy dictates the content of neural representations to an extent where individual word meanings are effectively masked, providing evidence that words do not have a one-to-one mapping in the brain. By showing that not even basic word meanings are impermeable to linguistic structure, our study highlights the inherent tension arising from combining stable concepts during real-time language processing.

## INTRODUCTION

Language allows us to string words together into endlessly creative linguistic expressions, like children assembling fantastical LEGO-brick structures. However, the analogy quickly breaks down: While identifying the yellow brick in a red LEGO tower is easy, there is (hopefully) no dog in *dog food* nor necessarily any cat in a *cat bed* (Figure 1). Unlike a LEGO brick, the meaning of a word is not fixed but shaped by its position in a structure, making language processing beyond the level of individual words a non-trivial task. How does the brain modulate a word’s meaning based on the construction in which it appears? Here, we directly examine how linguistic structure impacts word meaning by tracking the neural encoding of animacy across reversible noun-noun phrases. In doing so, we approach linguistic composition in the brain from a new angle, positioning our work between cognitive neuroscience research on the semantic system (Patterson et al., 2007; Binder et al., 2009; Huth et al., 2016; Ralph et al., 2017) and combinatory operations (Friederici et al., 2000; Bemis & Pylkkänen, 2011; Pallier et al., 2011; Zaccarella et al., 2015; Matchin et al., 2019).

**Figure 1:**
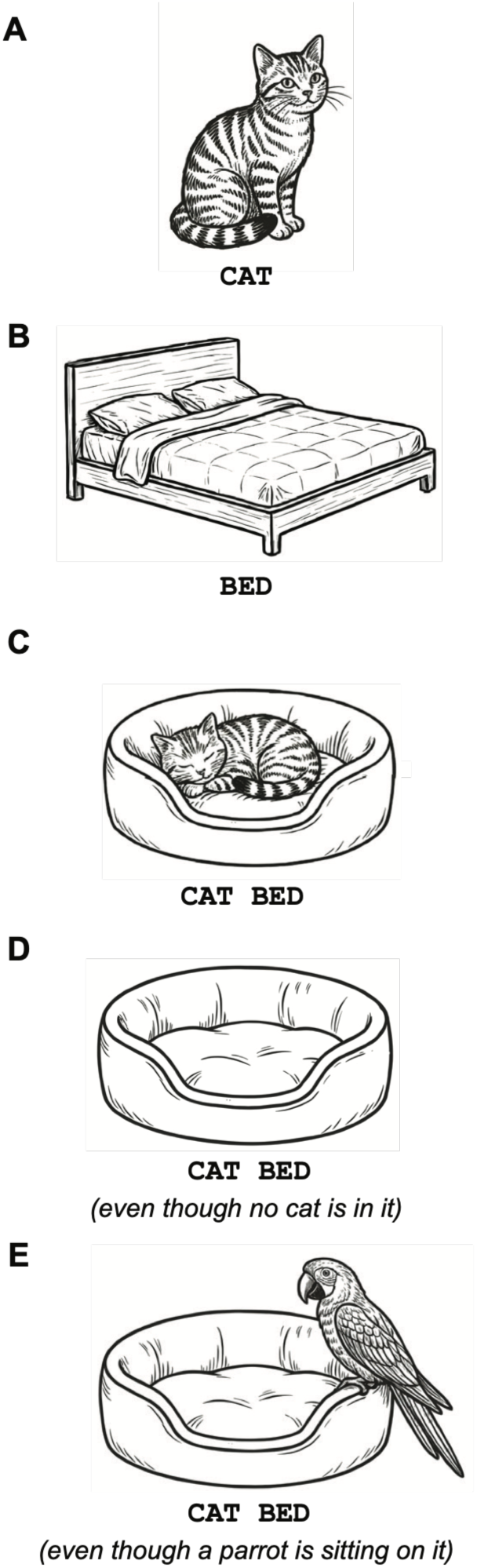
Word meanings are shaped by structural positions. Individual words compose together to form complex meanings. Take the words *cat* (**A**) and *bed* (**B**). Combining the two words as *cat bed* makes it a bed specifically for cats (**C**). We know, however, that a *cat bed* does not need to have a cat in it to qualify as such (**D**). In fact, *cat bed* remains a cat bed even when occupied by a parrot (**E**). While *cat* and *bed* thus both contribute to the concept *cat bed*, their relative importance in the composed concept varies as a function of their position in the linguistic structure.

Animacy is a fundamental semantic feature that organizes conceptual knowledge from an early age (Hatano & Inagaki, 1994; Rakison & Poulin-Dubois, 2001; Opfer & Gelman, 2011), allowing humans to separate animates (animals, including humans) from inanimates (non-living entities). The categorical nature of the animate-inanimate distinction is evidenced by studies of brain-damaged patients—often involving the anterior temporal lobes (ATL)—where access to animates or inanimates can be selectively impaired (Warrington & Shallice, 1984; Caramazza & Shelton, 1998; Henderson et al., 2021). Cognitive neuroscience provides independent support for an animacy-based partition of conceptual knowledge: Categories of animates and inanimates have distinct neural sig-natures when presented as pictures (Simanova et al., 2010; Murphy et al., 2011) or words (Simanova et al., 2010; Chan et al., 2011; Sudre et al., 2012), with parts of these representations overlapping across modalities (Leonardelli et al., 2019; Dirani & Pylkkänen, 2023). In noun-noun compounds, the brain even tracks the number of animate vs. inanimate features (Czypionka et al., 2023). While the importance of animacy thus only deepens the puzzle of why *dog food* need not involve a dog nor *cat bed* a cat, we can take advantage of the animate-inanimate distinction to track how word meanings are encoded in the brain.

Intuitively, some words in a phrase or sentence carry more weight than others— for example, in *dog food* and *cat bed,* the core meanings are conveyed by *food* and *bed*, not by *dog* and *cat.* In more formal terms, such modifier-head structures are interpreted asymmetrically, with phrase meanings being anchored by the head noun and the modifier interpreted only relative to that context (Kamp & Partee, 1995). In language processing, this could mean that not all semantic features of a word are activated when a word is encountered, with the animacy features of *dog* and *cat* remaining inactive in our examples because, as modifiers, their features are not features of the overall phrase meaning. This poses the question of its neural implementation: Do our brains commit to semantic features word-by-word, or do we wait until all evidence for the representation has unfolded? Clinical studies (Stark & Stark, 1990; Semenza et al., 2011; Marelli et al., 2013) and computational modelling of behavioral data (Marelli et al., 2017; Günther & Marelli, 2023) support differential encoding of words in noun-noun phrases based on functional roles, yet we still lack clear corresponding neural evidence (Ciapparelli et al., 2025).

We used spatiotemporally resolved magnetoencephalography (MEG) measurements to determine how linguistic structure affects the animacy encoding of words like *tiger* and *valley* across novel, reversible noun-noun phrases like *valley tiger* and *tiger valley*. Our primary decoding analyses, corroborated by secondary univariate analyses, queried whether the animacy of single words is retained when those same words function as either modifiers or head nouns.

## METHODS

### Participants

Thirty-seven native speakers of English with normal or corrected-to-normal vision were recruited through NYU’s SONA system and word-of-mouth to complete the experimental protocol. Participants provided their informed written consent and were compensated for their time. The study was approved by the Institutional Review Board (IRB) ethics committee of New York University (approval number: IRB-FY2024-8720). Following the exclusion of eight participants (based on excessive movement, sleepiness, or reporting a panic attack), the final dataset consisted of 29 participants (16 women, 9 men, 4 nonbinary; 20-40 years old, mean age ± SD: 26.2 years ± 5.1 years).

### Experimental design

To investigate whether linguistic composition impacts the neural signatures of word meaning, we employed a cross-condition decoding approach designed to test representational invariance. This involved training classifiers on neural data elicited by individual words and evaluating whether the learned representations generalized to words functioning as modifiers and head nouns in phrases. We had two distinct single-word training sets using (1) the same animals and locations functioning as modifiers and head nouns in phrases (ANIMAL/LOCATION) and (2) various animates and inanimates (ANIMATE/IN-ANIMATE). This two-pronged approach allowed us to adjudicate between the neural encoding of specific semantic categories vs. a more abstract notion of animacy.

#### Phrases

We created our phrasal stimuli by pairing 25 animals and 25 locations matched for lexical characteristics (e.g., length, frequency, orthographic and phonological neighbors; Balota et al., 2007; Brysbaert & New, 2009) in two plausible combinations (e.g., *tiger* with *valley* and *desert*) and presented each pair in both word orders (*valley tiger; tiger valley*). This yielded 50 animal-location pairs and 50 location-animal pairs. We chose reversible noun-noun phrases to keep lexical semantics constant (Graves et al., 2010), and we specifically opted for novel pairings (transition probability: 0% in COCA (Davies, 2008); < 0.07% in iWeb (Davies, 2018)) rather than existing ones like *house dog/dog house* to avoid confounding effects of word frequency and asymmetrical prediction effects across the two word orders. This low baseline predictability was corroborated by GPT-2 (small, 124M parameters) surprisal values for the second word given the first, which were high in both word orders (animal-location: mean = 16.36 bits, SD = 2.03 bits; location-animal: mean = 16.08 bits, SD = 2.61 bits), corresponding to extremely low probabilities. A paired *t* test indicated that the difference between the two word orders was not statistically significant, *t*(49) = 0.58, *p* = 0.56, *d*_z_ = 0.08, mean difference = 0.28 bits, 95% CI [–0.69, 1.26].

As part of a larger 2 × 2 × 2 design addressing distinct research questions, each pair was further modified by a real-world constrained geographical modifier (e.g., *Arabian valley tiger* rather than *Swedish valley tiger*) and a length-matched material modifier (*plastic*), and ungrammatical counterparts to the four grammatical phrase conditions were formed by swapping modifier ordering (*Arabian valley tiger; valley Arabian tiger)*. This paper focuses exclusively on the two grammatical conditions with geographical modifiers (i.e., *Arabian valley tiger* vs. *Arabian tiger valley*. See Krogh & Pylkkänen (2026a, 2026b) for discussion of the other experimental manipulations. All 400 trials (50 per condition) were presented in all-caps to mitigate the conventional capitalization of geographical modifiers.

To track word-by-word dynamics, we presented phrases using Rapid Serial Visual Presentation (RSVP; see Figure 2B). Following a fixation cross (200 ms) and a blank screen (200 ms), each word in the three-word target stimulus was presented in a 300 ms on, 500 ms off-sequence. A trial concluded with a 300 ms three-word task stimulus that was either identical to (match trial) or differed by one length-matched word from (mismatch trial) the target stimulus. Although this matching task can be performed via shallow processing, it reliably exhibits sensitivity to linguistic properties including, crucially, semantic content (Supplementary Materials of Fallon & Pylkkänen (2024)). For each participant, a trial was randomly designated as a match or a mismatch trial independently of experimental condition. All trials concluded with interstimulus intervals jittered between 600 and 750 ms.

**Figure 2:**
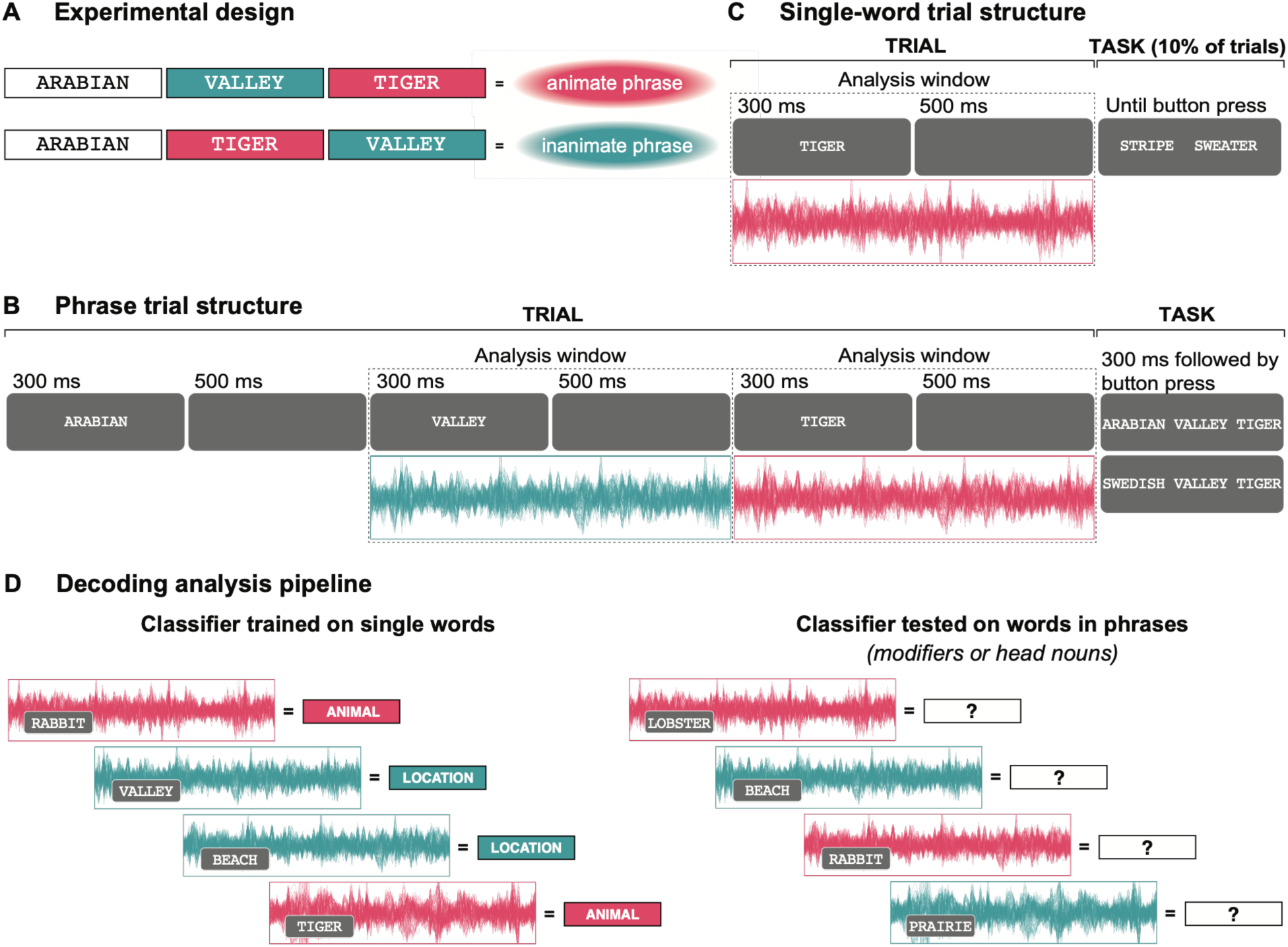
Experimental design, trial structures, and decoding analysis pipeline. (**A**) To investigate how linguistic structure shapes word meaning, our experimental design contrasted three-word phrases consisting of animal-location pairs in both word orders. Despite conflicting word-level animacy features, all stimuli were unambiguously animate or inanimate at the phrase-level. (**B**) The three-word phrases were presented word-by-word during continuous MEG recordings and followed by a matching task. The MEG analysis focused on neural data associated with nominal modifiers and head nouns (second and third word of a phrase). (**C**) Single-word trials functioned as a decontextualized base-line for animacy encoding. Participants saw two sets of single words, comprising either the same items used in the phrasal stimuli (ANIMAL/LOCATION) or a mixed group of nouns to test for abstract animacy (ANIMATE/INANIMATE). Each word was presented individually, also during continuous MEG recordings, with a semantic association task following 10% of trials. (**D**) Basic decoding analysis pipeline (for both temporal generalization and spatial decoding analyses). Decoders were trained to discriminate between single-word neural data with associated category labels (e.g., ANIMAL vs. LOCATION) and subsequently evaluated for their ability to predict category labels for words functioning as either modifiers or head nouns.

#### Single words: Animals and locations

Single-word trials with the same 25 animals and 25 locations used in the phrases provided a baseline measure of animacy at the level of semantic categories (ANIMAL/LOCATION) in the absence of structural context. Trials began with a fixation cross (200 ms) and a blank screen (200 ms) followed by a single word (300 ms) and another blank screen (500 ms). On 10% of trials, participants did a forced-choice semantic association task (e.g., *tiger* ◊ *stripe* | *sweater*). We chose this task to promote semantic processing of the stimuli and thereby support more robust classification results. Ten repetitions of each exemplar across ten blocks resulted in 500 trials (250 trials per condition).

#### Single words: Animates and inanimates

To evaluate whether our findings represented an abstract notion of animacy or simply reflected the specific semantic categories of animals and locations, we created a second training set of 25 animate and 25 inanimate words (ANIMATE/INANIMATE) that did not include animals or locations. Inanimates were drawn from the semantic categories of tools, vehicles, buildings, musical instruments, and pieces of clothing. Animates all referred to human or human-like individuals, ranging from relatively generic words (e.g., *tourist*) to words referencing nobility (e.g., *emperor*), professions (e.g., *dentist*), or supernatural creatures (e.g., *vampire*). Ideally, both sets should have spanned equally broad ranges of semantic categories, but the very nature of what it means to be animate—a feature unambiguously found only in humans and animals—precluded this. Stimulus selection criteria as well as trial and block structures otherwise mirrored those described in *Single words: Animals and locations.* Given the risk of participant fatigue, this experimental component was made optional, with 21 of the 29 participants opting to complete it.

### Procedure

The study took place at the NYU/KIT MEG LAB at New York University, New York. After providing informed consent and completing a brief demographic questionnaire, participants’ head shapes were digitized with a Polhemus FastSCAN system (Polhemus, VT, USA) to enable coregistration of the MEG data with the FreeSurfer *fsaverage* brain (Fischl, 2012). Digitizations included five future marker coil placements and the locations of three fiducial landmarks (nasion, left tragus, right tragus). Participants were given instructions about the structure of the experiment and familiarized with its components through a short set of practice trials. They were then set up in the magnetically shielded room hosting the MEG machine and completed the experiment in supine position during continuous MEG recording.

An experimental session presented 18 blocks in a semi-structured, pseudorandomized order that interleaved blocks from three experimental components: single-word ANIMAL/LOCATION trials (10 blocks of 50 trials), RSVP phrase trials (4 blocks of 100 trials), and a component briefly flashing the same phrase trials in full rather than presenting them serially (4 blocks of 100 trials). We included this third component to assess cross-presentation mode generalizability of combinatory processes; these findings are discussed in Krogh & Pylkkänen (2026a). Since the same phrases appeared in both presentation modes, appropriate counterbalancing was employed. Time permitting, participants were invited to do the optional component with single-word ANIMATE/INANIMATE trials (10 blocks of 50 trials), which they received additional compensation for. This was entirely voluntary as the core experimental protocol was already quite long. Participants took self-timed breaks throughout the recording session, which lasted on average 70 minutes without the optional component and 90 minutes with it.

### MEG data collection and preprocessing

A whole-head, 157-channel axial gradiometer system (Kanazawa Institute of Technology, Kanazawa, Japan) was used to acquire MEG data at a sampling rate of 1000 Hz and an online band-pass filter of 0.1–200 Hz. Marker coils were used to record participants’ head positions relative to the MEG sensors at the beginning and end of the recording session, and stimulus-to-trigger delays were measured and later corrected using a photodiode. As a first preprocessing step, the MEG data were cleaned of environmental noise with the Continuously Adjusted Least-Squares Method (Adachi et al., 2001) as implemented in the MEG160 software based on three reference channels. All subsequent preprocessing steps were conducted with MNE-Python (v.1.10; Gramfort et al., 2014) in Python.

The data were band-pass filtered at 1–40 Hz (the high-pass filter was necessary to attenuate NYC environmental noise) and flatlined or excessively noisy channels were removed and interpolated using data from neighboring channels. This included two known broken channels plus recording-specific channels identified via visual inspection (mean: 1.45 additional bad channels). Then, Independent Component Analysis (ICA) was used to remove well-characterized biological and environmental artifacts. Phrase trials were segmented into separate 800 ms epochs for modifiers and head nouns (300 ms word presentation; 500 ms blank screen) and baseline corrected with the 100 ms preceding the first word. Single-word trials were segmented into one 800 ms epoch (300 ms word presentation; 500 ms blank screen) with a 100 ms pre-stimulus baseline. Epochs were automatically rejected based on a 3000fT peak-to-peak amplitude threshold and manually excluded if trials were flagged during data acquisition due to technical issues or task-irrelevant behavior (e.g., visible sleepiness or talking to the experimenter). For phrase epochs specifically, we further removed trials with incorrect responses or response times more than three SDs from the participant mean. This yielded an average of 45.2 epochs (SD = 4.7) for each of the two phrase conditions analyzed here. For single-word trials, we similarly removed trials with incorrect responses as well as those immediately following the semantic association task (which occurred on only 10% of trials) to avoid potential spillover effects of task-induced surprisal. This resulted in an average of 222.6 epochs (SD = 5.9) per condition for the ANIMAL/LOCATION single-word trials and an average of 223.4 epochs (SD = 4.3) per condition for the ANIMATE/INANIMATE single-word trials.

Source estimation was done twice, first using single trial data for the spatial decoding analyses and then using balanced, condition-averaged data for the univariate analyses. First, we scaled and coregistered the FreeSurfer *fsaverage* brain to each participant’s digitized headshape and fiducials; this was used to generate a source-space mesh of 2,562 vertices per hemisphere (ico-4 spacing). Next, a forward solution was calculated using the Boundary Element Model (BEM) method, and channel noise covariance was estimated using the 100 ms pre-stimulus interval of all experimental trials. Together, the forward solution and covariance were used to calculate the inverse solution for each participant and applied to either single trials or condition-averages (SNR: 3), resulting in noise-normalized Dynamic Statistical Parameter Maps (dSPM; Dale et al., 2000).

Although the *fsaverage* brain was coregistered to individual headshapes, the use of a template in lieu of anatomical MRIs means that localization results should be interpreted with some caution.

### Statistical analysis

#### Behavioral data

The behavioral data cleaning largely mimicked that of the MEG data. We removed trials flagged during data acquisition and phrase trials with response times more than three SDs from the participant mean. For the single-word tasks, we simply report mean accuracies. For the phrase trials, we ran two (generalized) linear mixed effects regression models with phrasal animacy (animate, inanimate) and response type (match trial, mismatch trial) as fixed effects. Response type was included as a fixed effect since prior research has shown match and mismatch trials to incur different levels of processing difficulties (Fallon & Pylkkänen, 2024; Flower & Pylkkänen, 2024, 2026a; Krogh & Pylkkänen, 2025; Li & Pylkkänen, 2026). The dependent variables were either log-transformed response time data (cleaned of incorrect trials) or accuracy. The maximally converging random effects structure for reaction time had random intercepts for participant and item while the accuracy model also had a correlated by-participant random slope of phrasal animacy. Likelihood ratio tests were used to generate *p* values, and pairwise comparisons were subjected to a Tukey adjustment. Again, we restricted our analyses of the phrase trials to the two phrase conditions of interest for the present paper, though the results are qualitatively unchanged when using the full experimental design. All analyses were conducted using the *lme4* (Bates et al., 2015) and *afex* (Singmann et al., 2023) packages in R (v4.5.2) and RStudio (v2026.01).

#### Temporal decoding analyses with generalization across time

To identify the neural correlates of animacy, we performed time-resolved decoding with temporal generalization in sensor space (King & Dehaene, 2014). This involved training unique classifiers to discriminate between neural responses associated with different animacy categories and testing the classifier’s accuracy on unseen data. By moving beyond the temporal diagonal—rather than restricting training and testing to identical time points—we allowed for the emergence of clusters asynchronous across train and test times, accounting for the possibility that animacy representations may vary across different compositional contexts. Our four primary analyses involved training on one of the single-word sets (ANIMAL/LOCATION; ANIMATE/INANIMATE) and testing on the modifier or head noun in phrase trials.

Prior to analysis, the MEG data were downsampled to 200 Hz by averaging non-overlapping bins of 5 ms and reduced to 70 principal components (minimum variance explained across all analyses: 96.6%). This PCA mapping was learned from the training data and subsequently applied to the test data. The PCA components were scaled to unit variance and used as features for an l2-regularized logistic regression model. Regularization strength (C) was optimized based on a grid search on a logarithmic scale (10^-4^ to 10^4^) using stratified 5-fold cross-validation on the training set. We obtained temporal generalization matrices by training separate classifiers on pairs of time points in 100 ms windows (with edge padding), sliding in 10 ms increments across the 0–800 ms epoch. This entire pipeline was done separately for each analysis and each participant.

#### Group-level statistical testing of temporal decoding analyses

To assess above-chance decoding at the group-level, we employed non-parametric cluster-based permutation tests (Maris & Oostenveld, 2007) using a standard approach. We evaluated accuracy against a chance level of 0.5 using one-tailed, one-sample *t* tests at each train-test time pair and aggregated contiguous *t* values exceeding a *p* < 0.05 threshold into clusters. Cluster-level significance was determined by comparing the observed cluster statistics against surrogate distributions generated by shuffling condition labels within-subject (10,000 permutations). Corrected *p* values thus reflect the frequency of permuted cluster statistics that exceeded the maximum observed cluster statistic. Only clusters with a size of ≥ 2 points (equal to a minimum duration of 20 ms) were retained. Cluster boundaries should be treated as approximations as they are derived from uncorrected *t* values (Sassenhagen & Draschkow, 2019).

We report results from two analysis windows: a targeted 100–500 ms window intended to capture established animacy encoding effects and a broader 100–700 ms window. This latter interval represents the maximum viable temporal range from our 0–800 ms epoch, given the 100 ms edge padding of the sliding window architecture. The decoding analyses were performed in Python using *MNE-Python* (v1.0.3) and *scikit-learn* (v1.6.1; Pedregosa et al., 2011).

#### Spatial decoding analyses

Spatial decoding was performed in source space with a 10mm-radius searchlight. The procedure followed the temporal generalization analyses with two key distinctions: (1) Classification was performed at each individual neural source, and (2) with time samples within a specified window—rather than PCA components derived from sensor activity— as the neural features. Based on the temporal generalization results at the head noun, we performed these analyses in 50 ms × 50 ms train-test tiles across significant temporal clusters. To ensure that our spatial analysis focused on regions overlapping considerably with temporal clusters, we restricted our analysis to tiles with at least 20% coverage of the temporal cluster. Following the assumption that animacy, like other semantic features, is encoded as a spatially distributed pattern (Martin, 2007; Huth et al., 2016; Coutanche et al., 2020), we analyzed each hemisphere in its entirety.

Group-level significance in a temporal tile was assessed independently for each hemisphere, applying cluster-based permutation tests as described in *Group-level statistical testing of temporal decoding analyses.* Given that our spatial decoding analyses probe significant temporal clusters, they are to be considered post-hoc tests and the associated *p* values should therefore be interpreted as descriptive of spatial distribution rather than as independent inferential statistics. Furthermore, because decoding relies on distributed neural patterns, individual sources may be essential for classification without necessarily reaching significance when analyzed in isolation. As such, our spatial decoding results may only represent a subset of neural sources contributing to the temporal decoding results.

#### Univariate source-localized analyses

To complement our decoding analyses, we performed three spatiotemporal, cluster-based permutation tests on condition-averaged source time course data. First, a 2 × 2 repeated-measures ANOVA was conducted on the phrase trials with factors word position (modifier, head noun) and phrasal animacy (animate, inanimate). We specifically looked for interaction effects to elucidate how word position modulates the representation of animacy, using planned pairwise temporal clustering tests within the spatiotemporal cluster-extent to determine a cluster’s functional properties. Second, paired *t* tests were performed on each of the two single-word sets to contrast animacy (ANIMAL/LOCATION; ANIMATE/INANIMATE). All analyses were run individually in each hemisphere in a targeted (100–500 ms) and a maximally exploratory (0–800 ms) window, with the latter simply reproducing the clusters of the former. The cluster-based permutation tests were performed as described under *Group-level statistical testing of temporal decoding analyses,* with the exception that cluster contiguity spanned both time (≥ 10 ms) and space (≥ 20 sources). All spatiotemporal, cluster-based permutation tests were performed in Eelbrain v.0.40.4 (Brodbeck et al., 2023).

## RESULTS

### Distributed neural patterns encode animacy of head nouns, not modifiers, in the right anterior temporal lobe

We first took advantage of the millisecond temporal resolution of MEG and assessed, at each time point, classifier performance when trained on the sensor-level spatial neural patterns of single-word trials (ANIMAL/LOCATION; ANIMATE/INANIMATE) and tested on the sensor-level spatial neural patterns of modifiers or head nouns in phrases (Figure 2D). Crucially, the use of single-word trials as training data ensured that any successful decoding words in phrases cannot be attributed specifically to position-related differences in predictability. Since animacy may be encoded at different latencies when a word is presented in isolation and as part of a phrase, our temporal decoding approach employed temporal generalization (King & Dehaene, 2014). This involved training and testing the classifier across all pairs of time points rather than restricting the analysis to the same train-test times.

Our temporal decoders successfully discriminated between words like *tiger* and *valley* functioning as head nouns, but not when those same words functioned as modifiers (Figure 3). Specifically, the classifier trained on single-word ANIMAL/LOCATION trials and tested on head nouns yielded a significant diagonal cluster (train times: ∼220–380 ms; test times: ∼210–400 ms; *p* = 0.012), showing that animacy was encoded at similar latencies across the training and test data. In contrast, training on single-word ANIMATE/INANIMATE trials and testing on head nouns yielded an off-diagonal cluster, with delayed train times (∼290–440 ms) relative to test times (∼190–340 ms; *p* = 0.017). We did not observe any significant clusters when testing classifiers on modifiers (ANIMAL/LOCATION training: lowest *p* = 0.271; ANIMATE/INANIMATE training: lowest *p* = 0.152). The timings of the significant clusters conform well with the canonical N400 response, indicating that word meanings are retrieved by ∼300–400 ms (Kutas & Hillyard, 1980). The clusters also fall within the ∼100–600 ms window identified by prior decoding studies as capturing the animate-inanimate distinction (Simanova et al., 2010; Chan et al., 2011; Sudre et al., 2012; Leonardelli et al., 2019; Dirani & Pylkkänen, 2023). That cluster test times converged across training sets (ANIMAL/LOCATION; ANIMATE/INANIMATE) suggests that our decoders captured an abstract notion of animacy rather than one tied to semantic categories. This abstraction may also explain the asynchronous train-test times observed for ANIMATE/INANIMATE training: Extracting the animacy feature from a diverse set of words is likely more demanding, and therefore temporally delayed, compared to the processing of semantically more homogeneous categories like animals and locations.

**Figure 3:**
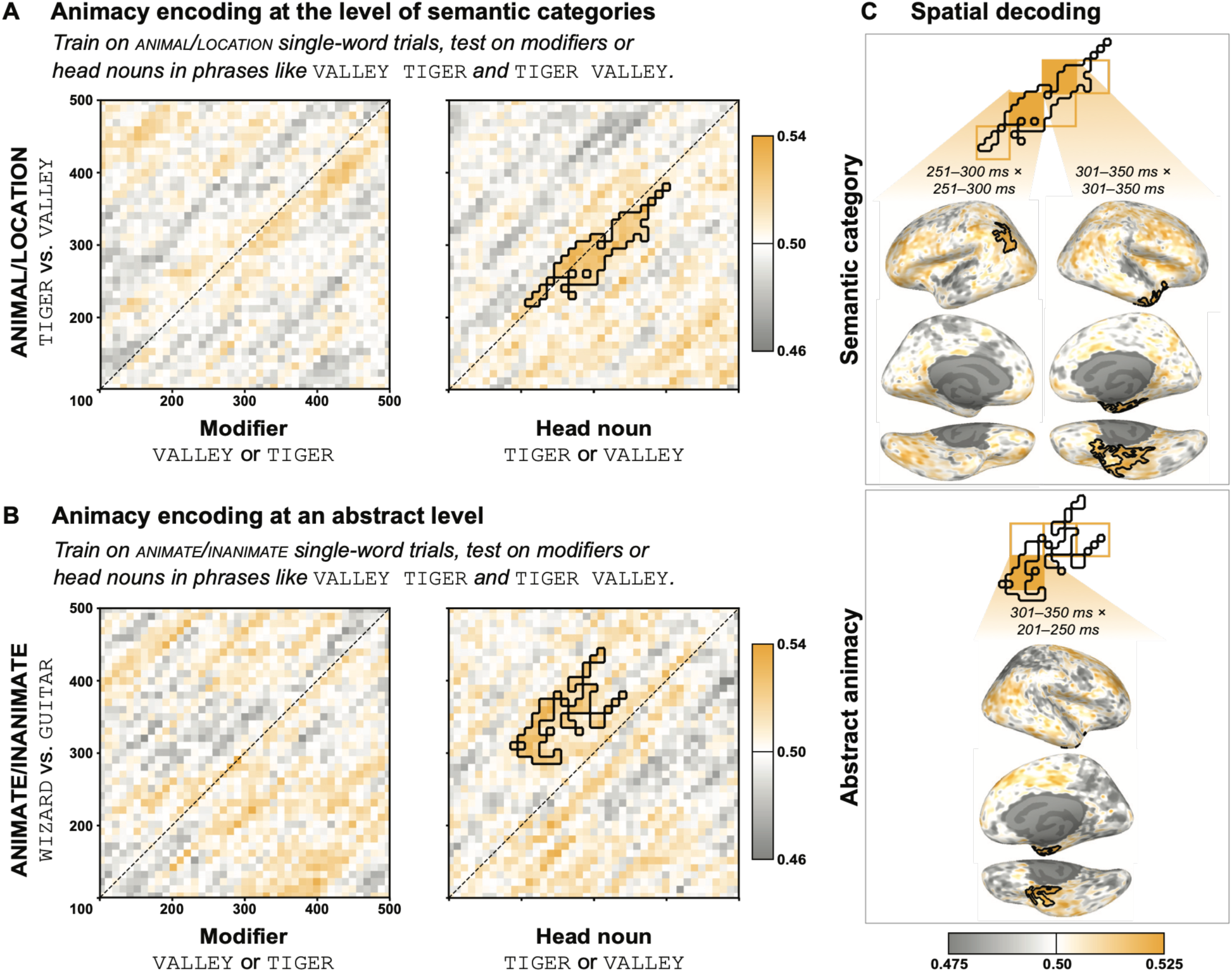
Animacy encoding in the brain across time and functional roles. Distinct classifiers were trained on single-word trials—either specific semantic categories (ANIMAL/LOCATION; **A**) or abstract animacy (ANIMATE/INANIMATE; **B**)—and tested on their ability to predict the category labels of modifiers or head nouns within phrases such as *valley tiger* and *tiger valley*. Matrices represent classifier train and test times (in ms), color bars indicate classifier accuracy. Significant decoding was observed only at the head noun, despite identical lexical contrasts at the modifier. This suggests that the brain does not represent animacy incrementally if later information can change the feature. While training on ANIMAL/LOCATION trials yielded a diagonal temporal cluster, ANIMATE/INANIMATE training resulted in a temporal shift, suggesting that is it more demanding to extract an abstract feature from a diverse group of words than words belonging to a single semantic category. Post-hoc spatial decoding analyses revealed involvement of the right anterior temporal lobe (ATL) regardless of training set (**C**) and involvement of the left parietal lobe when training on single-word ANIMAL/LOCATION trials. To capture the full extent of cluster activity, the significant clusters of the temporal decoding analyses were partitioned into smaller train-test tiles and evaluated independently.

Although temporal decoding relies on distributed neural patterns, certain brain regions may contribute more to these patterns than others. We therefore performed targeted spatial decoding analyses to localize the temporal decoding effects. Following our original cross-decoding procedure, we simply inverted time and space: Whereas temporal decoders were trained and tested on sensor-level spatial neural patterns at pairs of time points, spatial decoders were trained and tested on temporal neural patterns localized to each neural source (for similar approaches, see e.g., Gwilliams & King (2020), Zuanazzi et al. (2024)). To capture full cluster activity and facilitate comparison between on- and off-diagonal effects, we partitioned each temporal decoding cluster into 50 ms × 50 ms train-test tiles for separate spatial decoding analyses. Since these spatial decoding analyses targeted effects already identified in the temporal decoding analyses, they function as post-hoc tests, with the resulting *p* values characterizing the statistical strength of localized differences rather than serving as independent inferential statistics. ANIMAL/LOCATION training resulted in a left parietal cluster in the 251–300 ms × 251–300 ms tile (*p* = 0.036) and a right ATL cluster in the 301–350 ms × 301–350 ms tile (*p* = 0.001; Figure 3C). A similar right ATL cluster emerged with ANIMATE/INANIMATE training, albeit in an earlier test window (301–350 ms × 201–250 ms tile; *p* = 0.046). While spatial decoding localizations may not capture effects relying on broad, population-level representations across cortex, the consistent recruitment of the right ATL across training sets is striking.

### Increased activation for single-word animates and all animate-phrase constituents in the left hemisphere

Beyond representational similarity, the impact of structure on word meaning can be investigated by testing whether the same words evoke dissociable neural signals based on their functional roles. We therefore performed univariate, spatiotemporal analyses with signal strength rather than neural patterns as the dependent measure. First, we tested phrase-related neural signals for interaction effects between phrasal animacy (animate, inanimate) and functional role (modifier, head noun). Our analysis yielded one cluster from ∼288–455 ms (*p* = 0.015; Figure 4A), with increased amplitudes for words in animate phrases across a left antero-medial distribution encompassing combinatory regions like the left ATL (Bemis & Pylkkänen, 2011) and Broca’s area (Pallier et al., 2011). Sources within the cluster did indeed show activity differences for animals when functioning as modifiers vs. head nouns (323–370 ms; *p* = 0.001), with a similarly timed, non-significant activity divergence observed for locations (332–349 ms; *p* = 0.112). Activity was never significantly different between modifiers and head nouns *within* a given phrase (modifier vs. head noun in animate phrases: no clusters found; modifier vs. head noun in inanimate phrases: 440-452 ms, *p* = 0.1). Together, this supports our temporal decoding results: Structure modulates the neural representations of words.

**Figure 4:**
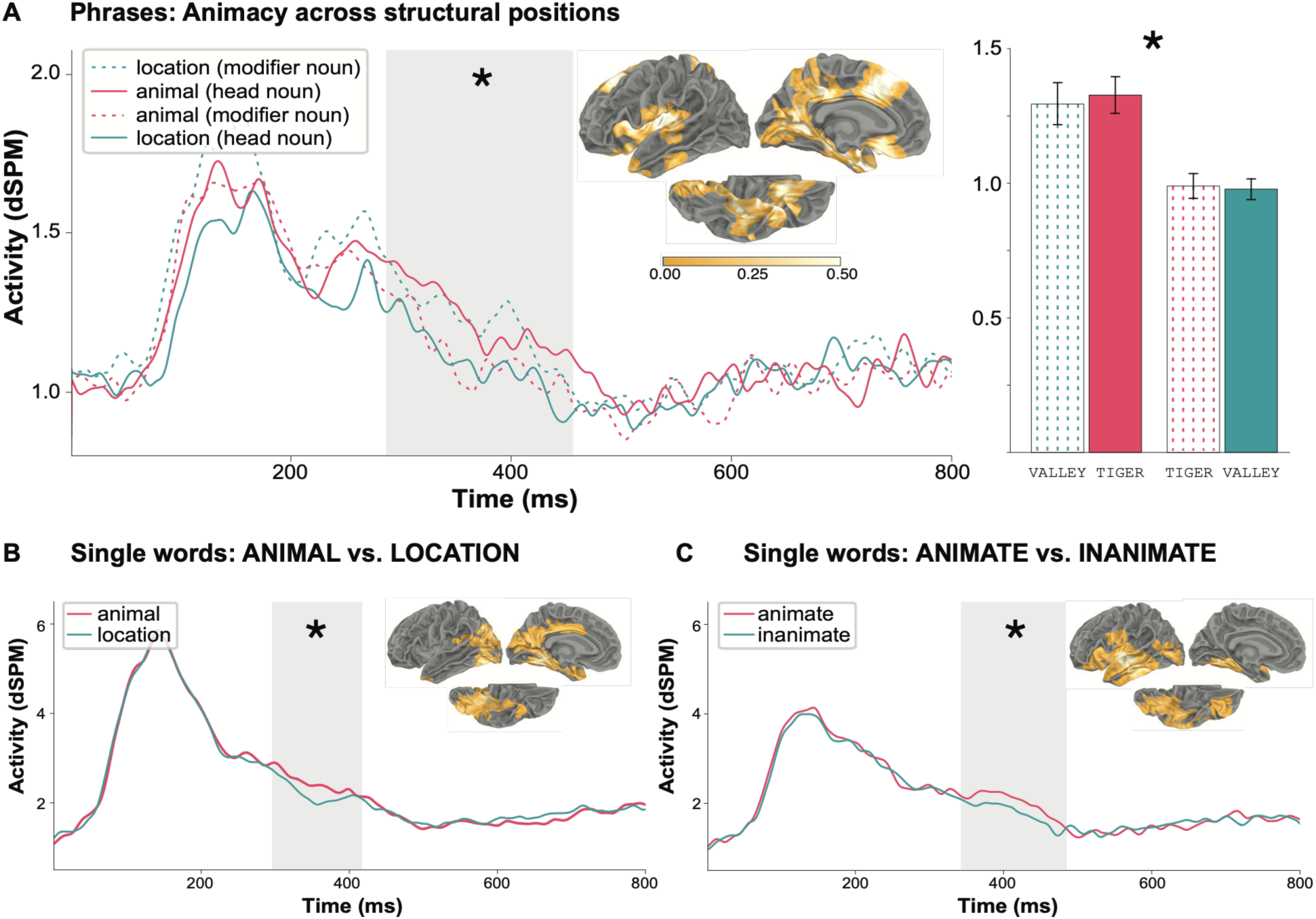
Source-localized results of univariate analyses. **(A)** Although our temporal generalization analyses only detected evidence of animacy encoding at the head noun, univariate analyses revealed temporally similar activation patterns for nouns within the same phrase across the left hemisphere, with higher amplitudes for both constituents of animate phrases relative to those of inanimate phrases. **(B)** While animals and locations embedded in the same phrase were not statistically different from one another, they are clearly distinct as single words, with increased amplitudes for ANIMAL over LOCATION trials. (**C**) Similar activation increases are seen for ANIMATE over INANIMATE trials.

Although our phrase-level univariate cluster overlaps temporally with the decoding results, its left-hemisphere localization contrasts with the right ATL decoding localization. We therefore repeated our univariate analyses for the single-word sets to determine if combinatory operations drove this left-hemisphere effect. The left-lateral clusters obtained for the single-word analyses, however, dispel such an interpretation. Specifically, single-word ANIMAL/LOCATION trials resulted in a medial, temporo-occipital cluster from ∼297–416 ms (*p* = 0.031; Figure 4B) while single-word ANIMATE/INANIMATE trials produced a later, more lateral fronto-temporal cluster from ∼344–483 ms (*p* = 0.041; Figure 4C), with ANIMAL/ANIMATE eliciting more activation than LOCATION/INANIMATE.

### Shared univariate signatures for within-phrase semantic categories do not imply shared neural representations of animacy

The comparable within-phrase signal strengths for modifiers and head nouns in our phrase-level univariate results raise a question about their underlying source. Does this uniformity arise from anticipatory encoding of head noun animacy at the modifier, triggered by the predictive experimental context? If so, phrase-level animacy may have masked the modifier’s word-level animacy, preventing successful decoding. Alternatively, does this uniformity reflect the integration of full modifier meaning into the head noun, rendering *valley* in *tiger valley* distinct from a generic *valley*? While the animacy of head nouns was decodable, it remains possible that the neural patterns likewise contained traces of the modifier’s meaning.

We explored this question in post-hoc temporal decoding analyses. As before, we trained our decoders on single words and tested them on words in phrases, but this time we swapped the category labels of modifiers and head nouns. If head noun animacy is prematurely encoded, a decoder should achieve above-chance accuracy when classifying the modifier *tiger* as LOCATION/INANIMATE in *tiger valley*. Conversely, if the full modifier representation integrates with the head noun, a decoder should achieve above- chance accuracy when classifying the head noun *valley* as ANIMAL/ANIMATE in *tiger valley*. None of these post-hoc analyses returned clusters in the 290–455 ms window of the univariate analysis (all *p* > 0.234; Figure 5). However, single-word ANIMAL/LOCATION training did reveal late anticipatory encoding of head noun animacy at the modifier position in a maximally exploratory window (Figure S1; train times: 130–320 ms; test times: 580–700 ms, *p* = 0.047). The comparable within-phrase activation in our univariate analyses therefore cannot be attributed to one word adopting the full neural representation, including the animacy feature, of the other. Crucially, this null result does not preclude such shared activation from reflecting the integration of features beyond those captured by our animacy classifiers.

**Figure 5:**
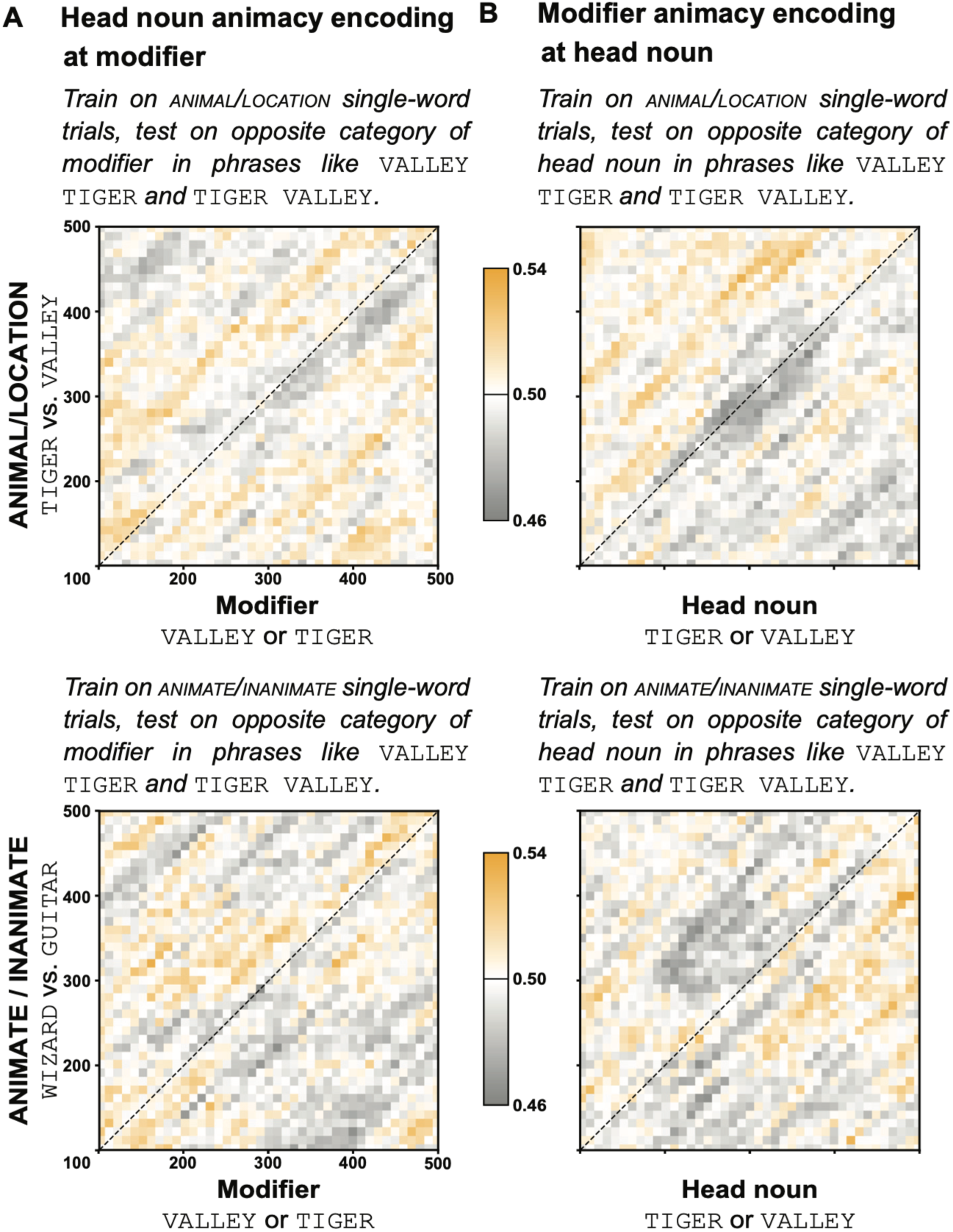
Post-hoc temporal decoding analyses swapping category labels. To investigate the comparable activation levels for words in the same phrase as revealed by the univariate analyses, post-hoc temporal decoding analyses probed whether head noun animacy is anticipatorily encoded in the modifier position (**A**) or if modifier animacy is integrated at the head noun (**B**). This was achieved by swapping the category labels associated with modifiers and head nouns. For example, anticipatory head noun encoding was tested by associating the label LOCATION/INANIMATE with *tiger* in *tiger valley*. None of these post-hoc analyses yielded results overlapping with the significant time window of the univariate analysis.

## DISCUSSION

### Structure shapes word meaning

By tracking the neural encoding of animacy in reversible noun-noun phrases, we investigated whether linguistic structure modulates word meaning. Is the meaning of *tiger* the same across *valley tiger* and *tiger valley*? Our results suggest not: While animacy as an abstract feature was decodable for head nouns from ∼200–350 ms, this distinction was statistically undetectable when those same words functioned as modifiers. Although previous research found that prior word representations endure at subsequent words (Fyshe et al., 2019), our findings suggest that word-level semantic features are suppressed if compositionally irrelevant, consistent with recent evidence that combinatorial context can substantially alter conceptual representations (Law et al., 2026).

What are the functional implications of words having distinct neural representations across different structural positions? Words in modifier positions are clearly not devoid of meaning—after all, we can conjure different concepts for novel noun-noun phrases like *valley tiger* and *tundra tiger,* just as we easily distinguish *desk lamp* from *floor lamp* and *hairbrush* from *toothbrush.* How such conceptual composition unfolds has been a major focus in the psychology of concepts (Smith et al., 1988; Murphy, 1988; Wisniewski, 1997), although no extant theory can account for the full range of possible conceptual relations. Crucially, only certain aspects of the modifier constrain the interpretation of the head noun: A lamp’s size is determined by its location, a brush’s bristle density by its use, and a tiger’s physical traits (e.g., fur thickness and pigmentation) by its habitat, whether lush valleys or icy tundra. In contrast, the inanimacy of *valley* is irrelevant (and, in fact, contradictory) for *valley tiger,* much as a desk having four legs is irrelevant for *desk lamp*. This suggests that the modifier’s neural signature reflects a representation pruned of unnecessary features to facilitate integration with the head noun, a finding consistent with behavioral work showing differential weighting of conceptual features (e.g., Barclay et al., 1974; Barsalou, 1982). While our animacy-based decoders focus on the presence of a specific semantic feature, the comparable within-phrase activation observed in the univariate analysis does suggest some process of feature integration across words.

This study demonstrates that the brain efficiently encodes word meanings based on their functional roles within fixed-length phrases. How might these findings translate to naturalistic discourse? While different from constrained laboratory settings, grammatical and contextual cues likewise make language processing “in the wild” highly predictable (Federmeier, 2007; Kuperberg & Jaeger, 2016; Pickering & Gambi, 2018; cf. also Nieuwland & Van Berkum (2006) on how discourse context can override compositional animacy expectations). Consider a sentence like *We have no more dog food.* The absence of a determiner rules out a singular count-noun reading while real-world knowledge blocks a mass-noun reading (as is possible with *chicken* when referring to meat), allowing us to immediately identify *dog* as a modifier. Similarly, in *Did Max rip apart the dog bed or the cat bed?,* structural parallelism allows featural pruning of the modifier *cat*. Extralinguistic context provides similar constraints; for example, a shopper searching for a replacement cat bed at the pet store would so strongly anticipate the head noun *bed* upon hearing *Can I help you find your new cat …* that the animacy of the modifier *cat* can be disregarded. Clearly, there is potential for our findings to generalize to more naturalistic settings. Whether such structure-dependent encoding of word meaning is learned on par with other statistical regularities found in language is an exciting question for future research.

### Hemispheric asymmetry: Lexical vs. abstract animacy in the anterior temporal lobes

Intriguingly, our results revealed a hemispheric dissociation in the encoding of animacy: Our decoding analyses implicated the right ATL while univariate effects localized to, among other regions, the left ATL. Crucially, because each hemisphere was evaluated using both approaches, such cross-method divergence is not an artifact of ROI selection. This asymmetry may therefore offer a new perspective on ATL specialization. Although both ATLs are critical neural substrates for semantic knowledge, meta-analyses of neuroimaging (Rice et al., 2015) and neuropsychological data (T. Lee et al., 2002) suggest a left-hemisphere bias for verbal memory and a potential right-hemisphere bias for nonverbal memory (T. M. C. Lee et al., 2002; Vaz, 2004). Our results align with this distinction in semantic storage. Specifically, left ATL univariate effects arose from direct word comparisons, thus relying on lexically grounded representations, whereas the feature extrapolation of the decoding analyses suggests that the right ATL effects tapped into a more abstract notion of animacy. Should these patterns generalize to other semantic features, this would provide a robust framework for understanding how the two hemispheres store conceptual knowledge.

## CONCLUDING REMARKS

In this study, we found that animacy—operationalized as semantic categories (ANIMAL/LOCATION) or at a more abstract level (ANIMATE/INANIMATE)—was neurally encoded for words used as head nouns (*tiger* in *valley tiger*), but not when those same words functioned as modifiers (*tiger* in *tiger valley*). That our brains represent *tiger* in *valley tiger* as an entirely different animal from *tiger* in *tiger valley* suggests that compositional processes actively reconfigure word meanings, prioritizing phrase-level meaning over word-level features.

While word meaning variability in context has been extensively studied using polysemes (e.g., Beretta et al. (2005); Pylkkänen et al. (2006); Klepousniotou et al. (2012); Messi & Pylkkanen (2025)) and homographs (e.g., C. Lee & Federmeier (2006, 2009)), our work distinguishes itself by investigating how a specific word meaning is modulated by its functional role. That a word can have fundamentally different neural instantiations based purely on structural context highlights the remarkable flexibility of the brain when “doing” language. Words are thus not just static entities retrieved from a mental lexicon but dynamically shaped by structural demands, raising exciting questions about the linkage between stable conceptual knowledge and the fluid representational states required for real-time language use.

## AUTHOR CONTRIBUTIONS

SK and LP designed research; SK performed research; SK analyzed data; SK wrote the paper; SK and LP edited the paper.

## ACKNOWLEDGEMENTS

This work was supported by the National Science Foundation award #2335767 (LP). We thank Bernarda Basualdo for assistance with data collection as well as Alec Marantz, Ailís Cournane, Sebastian Michelmann, Alona Fyshe, and the members of NeLLab for their suggestions and feedback. Cursor, an AI-assisted code editor, was used for limited code writing, editing, and debugging of existing analysis scripts, using both automatic model routing (Auto) and manually selected Claude Sonnet and Opus models (4.5 and 4.6). Google Gemini was used for minor editing in the final writing stages. All output was reviewed and edited by the authors, and the authors take full responsibility for the content of the published article.

## SUPPLEMENTARY MATERIALS FOR

### SUPPLEMENTARY TEXT

#### Behavioral results

Our statistical modelling of behavioral data associated with the matching task in phrase trials did not produce significant effects of animacy. While response times showed no modulation by animacy (*p* =0.215), accuracy demonstrated a marginal trend toward significance (*p* = 0.083), with higher accuracies for animate phrases (i.e., phrases like *valley tiger*). Match trials were processed significantly faster and more accurately than mismatch trials (both *p* < 0.002), aligning with prior studies using this matching task (Fallon & Pylkkänen, 2024; Flower & Pylkkänen, 2024, 2026; Krogh & Pylkkänen, 2025; Li & Pylkkänen, 2026). We did not see any significant interaction effects (both *p* > 0.516).

**Figure S1:**
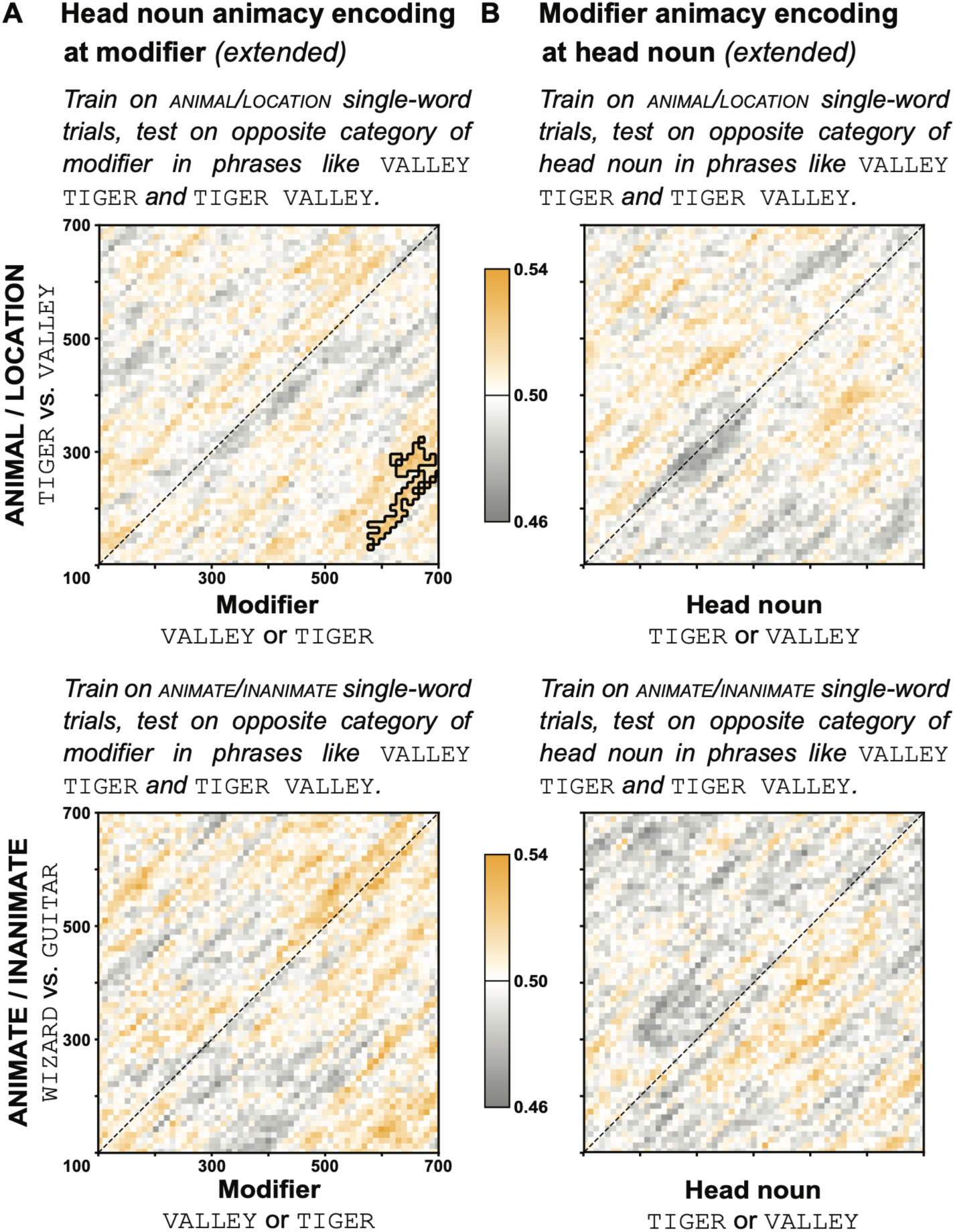
Post-hoc temporal decoding analyses swapping category labels (extended time window). Extending the time window from 100–500 ms to 100–700 ms in the post-hoc temporal decoding analyses yielded one cluster from the classifier trained on single-word ANIMAL/LOCATION trials, with delayed test times relative to train times (**A**; top). This suggests that while head noun animacy *is* prematurely encoded, such encoding happens only shortly before stimulus presentation. The extended time windows did not yield any evidence for modifier animacy being integrated at the head noun (**B**). Matrices represent classifier train and test times (in ms), color bars indicate classifier accuracy.

